# Pre-implantation genome-wide methylation enables environmental adaptation in a social meso-carnivore

**DOI:** 10.1101/2024.07.07.602321

**Authors:** Tin Hang Hung, Ming-shan Tsai, Chris Newman, David W. Macdonald, Christina D. Buesching

## Abstract

Many wild populations are increasingly stressed by rapid climatic change. While behavioural plasticity can enable limited tactical adaptive responses, standing genetic variation limits the species’ capacity to respond to climate change velocity. Epigenetic modification may provide a more rapid and plastic adaptive mechanism, but has been little studied in wild-living animals. Here we investigated CpG methylation during the pre-natal and early-life development of 95 European badger cubs between 2003 and 2011). During 10-months of delayed pre-implantation variability in precipitation between previous year’s February and April was the top determinant of methylation patterns among neonates, followed by mean temperature and temperature variability. Among the 4,641 significant weather-associated CpG sites, most occurred in the 47S rDNA region. Methylation of 47S rDNA was also associated with early-life weight, implying a mechanism that relays environmental stress to phenotypic stress. We also detected evidence for predictive adaptive response. Among the 1,641 CpG sites associated with early-life weight, pathways were associated with early-life growth, immune regulation, and to the development of aggression for competitive access to weather-limited food resources were over-represented. We conclude that a species’ epigenetics can have an important role in adaptive plasticity to environmental changes with important implications for biodiversity conservation and management.

## Introduction

In response to rapid climate change^1–3^ and environmental stressors, natural populations face three options: to migrate, adapt *in situ*, or suffer extirpation^4^. Migration is not always viable depending on the dispersal capacity of the species, the availability of habitats, niche competition^5^, and landscape barriers^6^. While behavioural plasticity can provide limited tactical benefits for species under certain scenarios^7,8^, enduring genetic adaptation is limited by the species’ capacity for variation in those traits under selection^9^. Furthermore, reliance on standing genetic variation^10,11^ (Haldanes’s sieve^12,13^) may not keep pace with climate change velocity^14,15^, particularly for populations with low gene flow rates and low genetic variation^16^. Rapid climate change typically exerts stronger selection pressure on early-life stages in most organisms, which are most vulnerable to environmental stress^17^, causing generally negative developmental effects on phenotypes^18^. In addition, extreme weather events can induce a shift in selection pressures over short timeframes within an individual’s life history^14,15^.

There has been growing appreciation of the evolutionary importance of epigenetic modification^19^ and how this could function as a rapid and plastic mechanism enabling species to adapt to changing environments^20–22^ and climatic conditions^19,23,24^. Epigenetic modifications affect the regulation of gene expression without changes to the DNA sequence involving the covalent binding of chemical groups to either DNA or histone proteins that are not erased by cell division and are thus heritable^25^, potentially overcoming the limits of standing genetic variation^26^.

Methylation of cytosine residues in a cytosine-phosphate-guanine (CpG) dinucleotide to form 5-methylcytosine (5mC) is the most well-established epigenetic modification in vertebrates^27,28^. 5mC regulates gene expression depending on its genomic location: promoter methylation results in stable silencing of gene expression^29^, while gene body methylation is associated with a higher level of gene expression only in dividing cells^30^. Mammalian methylomes undergo dynamic changes during early-life development^31^. *De novo* methylation occurs at the blastocyst stage during implantation, when the inner cell mass cells start to differentiate to form the embryonic ectoderm, facilitated by DNA methyltransferases DNMT3A/3B^32^.

Various studies have demonstrated epigenetic effects on early-life developmental programming in response to environmental stress^33–35^. Methylation of genes associated with growth (e.g. GRB10 and MSX1)^36^, stress responses (e.g., NR3C1)^37^, or cellular metabolic homeostasis (e.g. rDNA)^38^ cause distinct phenotypic differences in both human neonates and in non-human animal models. In wild-living mammals, climate change and extreme weather conditions can cause habitat degradation and loss^39^. In turn, this can disrupt foraging success, causing nutritional stress^40^ and lower resistance to pathogens^41^ among prospective parents, which can result in prenatal and early-life adversity in their offspring.

Despite the importance of understanding how environmental stressors can cause epigenetic effects in the early-life development of animals, little is known from wild populations (but see ^42,43^), limited by the availability of data covering a long enough period of variable weather conditions sufficient to detect any signal of adversity^42^ and the lack of complete genome information for most wildlife species. To address this knowledge gap, here we utilise multi-cohort data (1987 to 2019) from a systematically monitored population of European badgers (*Meles meles*; hereafter ‘badger’) at Wytham Woods (Oxfordshire, United Kingdom^44,45^), complemented by a recently assembled reference genome for this species^46^.

Badgers provide a highly informative model for investigating the responses of a generalist species to weather effects in a temperate region^47^. Rainfall and temperature extremes affect badger activity regimes^48^, bodyweight^49,50^, and life-history^51^. These effects operate via prey availability, primarily earthworms (*Lumbricus terrestris*) in our study region, which are sensitive to microclimate and weather conditions^50,51^, as well as the exposure of badgers to inclement weather when active^52^. Cubs are substantially more vulnerable to impoverished food supply and exposure to harsh weather than are adults^53,54^, compounded by severe coccidiosis (caused by *Eimeria melis*) that develops during immunological immaturity, resulting in malabsorption, steatorrhea, and dehydration^55,56^. Cubs are thus more susceptible to oxidative stress and oxidative damage than adults^57^. In turn, poor weather conditions during *in utero* and early-life development are associated with shorter early-life (3–12 months old) telomeres^58^ and higher rates of gamaherpesvirus reactivation ^59^. Importantly, inter-annual variation in weather conditions can exert substantially different effects between cohorts^60^. Badgers are also one of relatively few mammal species with delayed implantation^61^. This stretches from post-partum conception around mid-February until short day-length, modified by maternal body condition, triggers implantation in late December^62^. Thereafter, gestation lasts ∼56 days until a mean litter size of 1.5 ± 0.3 cubs are born in the following February^58,63,64^. These highly altricial cubs then remain underground within natal chambers until they are 8–10 weeks old^65,66^, at which time weaning commences.

Importantly, during this extended pre-implantation period, badger blastocysts, in common with other carnivores, remain metabolically active^61,67–69^, consuming oxygen and synthesising DNA, RNA, and proteins, although at slower rates than in activated blastocysts^70^. Furthermore, blastocysts continue to increase in size during pre-implantation, due to fluid accumulation within the blastocoele and active cell division in the trophoblast^70^, growing in diameter from ∼0.1 mm to up to ∼4.0 mm during pre-implantation^71^. It is, therefore, likely that any physiological stress affecting nutrient availability, oxygen availability, or redox state will affect blastocyst metabolic pathways, with deficits impairing implantation potential and subsequent foetal development. Harsh weather conditions post-conception cause females cumulative stress effects (a disease-focused effect) and environmental match–mismatch effects (an evolutionary-developmental effect)^72,73^. Therefore, blastocysts may potentially undergo methylation pre-implantation, arising through maternal effects, as well as during embryonic gestation and neonatal development^74,75^.

Here we sub-sampled a collection of whole blood samples^76^ from over 1,800 individual badgers^46^ to investigate for any link between weather conditions and methylation signals. Our objectives were: (1) to determine the most influential time window and weather metrics affecting the methylome during prenatal and early-life stages using a combined sliding window^77,78^ and machine learning^79^ approach; (2) to identify differentially methylated CpG sites in promoter regions and predict their functions using association analyses^80^; (3) to examine how methylation patterns are related to early-life weight; and (4) to assess whether these patterns reflect local spatial adaptation, using Procrustes residuals^81,82^.

## Methods

### Blood sample collection and preparation

Free-ranging badgers were captured in Wytham Woods, Oxfordshire, United Kingdom (∼51° 46’ 26” N, 1° 19’ 19” W) between 1987 and 2019, as part of a long-term badger research programme. Details of the study site and badger trapping and handling protocols can be found in ^83, 84^ and ^85^. Badger social group affiliation was inferred from trapping location. Individuals were (re-)identified from a serial tattoo number given at first capture^86^ which thereafter provided absolute age, except for a small minority of individuals for which age was inferred from toothwear^51^. Badgers were sedated^87^ and a variety of data were collected including each individual’s sex; whole blood was collected into EDTA-coated vacutainers via jugular venipuncture, then pipetted into microcentrifuge tunes and stored immediately at –20 °C. Cubs could not be caught and blood sampled until they were c. 12 weeks old because (i) they were initially underground, then subsequently they were (ii) still suckling and could not be separated from their mother, and (iii) too small to sustain anaesthesia safely^85^. For this study, we randomly selected 95 cubs, blood sampled at each individual’s first capture between late May and mid-June, representing cohorts born between 2003 and 2011 (except for 2006 when blood was not collected (**Supplementary Table 1**)).

### Library preparation and reduced representation bisulfite sequencing (RRBS)

Genomic DNA was extracted and purified from 200 μL of whole blood using Monarch Genomic DNA Purification Kits (New England Biolabs, United Kingdom). DNA quantity and quality were assessed using NanoDrop 2000 (Thermo Scientific, United States).

Samples of ∼200 ng genomic DNA were digested with 20 units of MspI (C↓CGG) and 10 units of AluI (AG↓CT) (New England Biolabs, United Kingdom) at 37°C overnight (∼16 hours). Digested DNA was purified using 1.8× AMPureXP magnetic beads (Agencourt, United States), then end-prepped and dA-tailed using the NEBNext Ultra II End Repair/dA-Tailing Module (New England Biolabs, United Kingdom). Methylated adaptors were ligated to end-repaired and dA-tailed DNA using the NEBNext Ultra II Ligation Module (New England Biolabs, United Kingdom), followed by USER enzymatic treatment. Ligated products were purified and size-selected for fragments shorter than 450 bp using 0.6× AMPureXP.

Size-selected fragments were bisulfite-converted using Methylamp One-Step DNA Modification Kit (Epigentek, United States) according to the manufacturer’s protocol. These sub-libraries were barcoded and enriched using NEBNext Q5U Master Mix (New England Biolabs, United Kingdom). Sub-libraries were purified using 1.8× AMPureXP and pooled to equimolarity for the final library at ∼20 nM, which was then sequenced on an Illumina NovaSeq 6000 system with a paired-end mode of 150 bp.

### Alignment of reads to reference genome and methylation calls

Raw reads were trimmed for adaptor contamination and low-quality bases (Phred score < 20) using Trim Galore 0.6.6. For adaptor-trimmed forward and reverse reads, an additional 2 base pairs were removed from 3’ and 5’ ends, respectively, to avoid artificial methylation calls due to filled-in cytosine positions close to the 3’ MspI digestion site. Reads were then aligned to the haplotype-resolved assembly for *M. meles*^46^ (mMelMel3.2; GCA_922984935.2) using bwa-meth 0.2.5^88^ (with bwa-mem2 2.2.1^89^). After sorting and indexing, duplicates in the resultant BAM files were marked using Picard 2.27.4^90^. Per-base CpG methylation level was extracted using MethylDackel. The output bedgraphs for all samples were collated to a unionbed using bedtools v2.30.0 and filtered for 25% missing data across samples. Remaining missing data were imputed using missForest 1.5.

### Sliding-window characterisation of weather metrics

Daily weather data for temperature (T) and precipitation (P) were obtained from the Radcliffe Meteorological Observatory Site in Oxford city (∼51° 45’ 22.79” N, 1° 15’ 30.00” W), within ∼5 km of the Wytham Woods study site. These data were then computed as mean monthly values and used as predictor variables to analyse how weather effects correlated with methylation patterns during pre- and post-natal development up until neonates reached three to four months old. For each badger cohort, we defined the post-fertilisation / pre-implantation (i.e., embryonic diapause^61^) period as occurring between February (conception) and November (earliest implantation) of the previous year (02_Y–1_ – 10_Y–1_), and the post-plantation / gestation period as occurring between December of the previous year and February of the birth year (11_Y–1_ – 02_Y+0_)^67^ (**Figure 1a**)^62^. Blood samples were taken around the end of May – early June of the birth year (05_Y+0_). This enabled us to test for any effects of methylation during early neonatal development over 16 preceding 1-month windows (e.g., 02_Y–1_ – 02_Y–1_, 03_Y–1_ – 03_Y–1_, …, 05_Y+0_ – 05_Y+0_), 15 2-month windows (e.g. 02_Y–1_ – 03_Y–1_, 03_Y–1_ – 04_Y–1_, …, 04_Y+0_ – 05_Y+0_), etc, resulting in a total of 16 + 15 + 14 + … + 1 = 136 potential time windows for each cohort.

**Figure 1.**
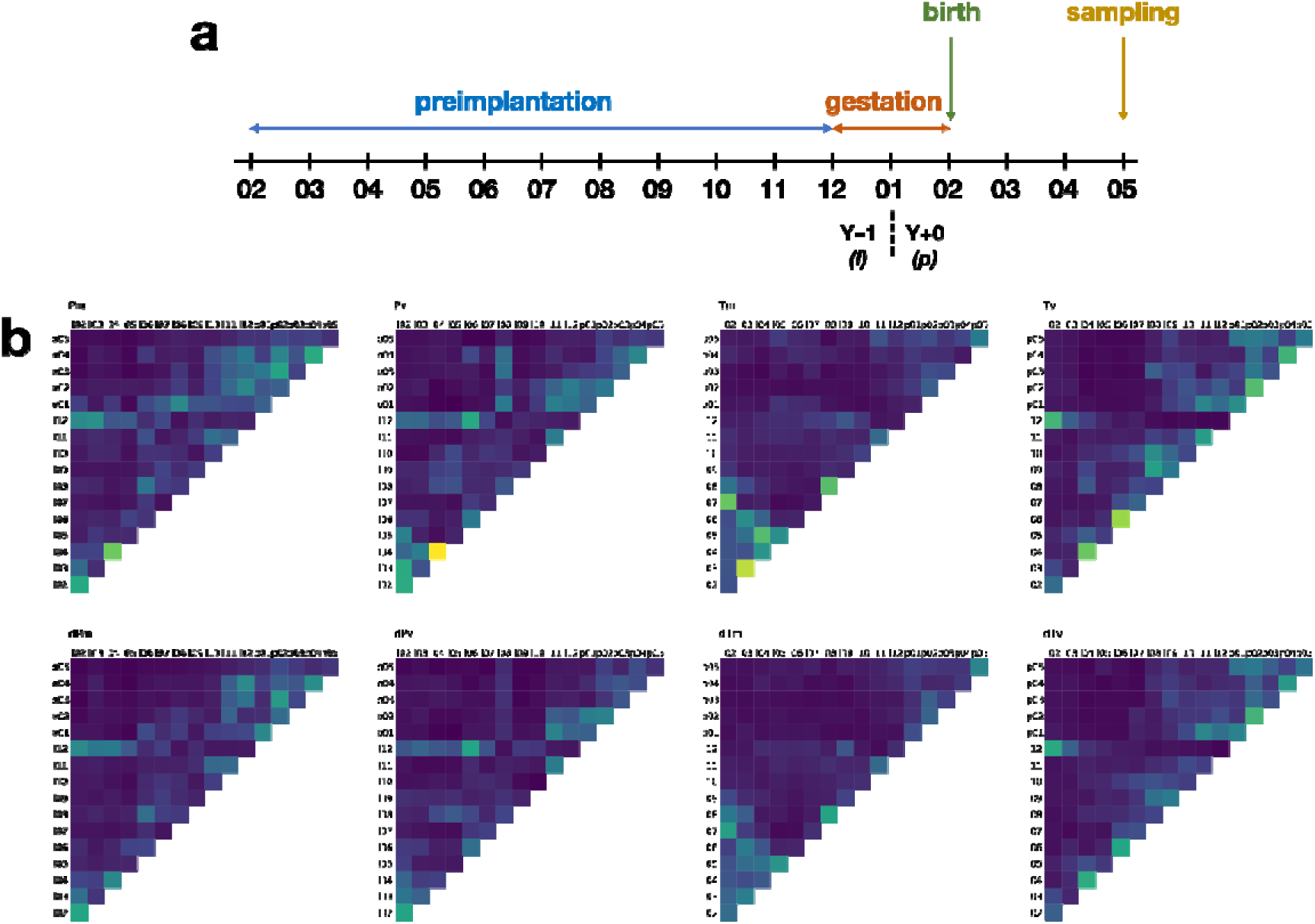
**(a)** Early life stages of preimplantation, gestation, and birth of a badger. **(b)** Heatmaps of *R*^2^-weighted importance of different sliding time windows for all environmental variables. The column axis denotes the start of the time window, while the row axis denotes the end of the time window. Thus, the diagonal of the heatmap shows the one-month time windows, while the top-left cell shows the full 16-month window from February of the previous year and the May of the current year.

We calculated four different statistics for Temperature (T) and Precipitation (P) across time windows, including the mean (m) and standard deviation within each window (v), the residual deviation for each month from the monthly mean for the five years prior and after the birth of the cohort (dm), and the deviation of monthly standard deviation from the same monthly time window over the five years prior and after the birth of the cohort (dv).

In summary, we characterised 1,088 weather metrics (= 2 parameters (temperature and precipitation) × 136 time windows × 4 statistics). These are denoted such that dTv_l07.p02 for 2015 represents the temperature standard deviation between July 2014 and February 2015 compared to the standard deviation for the same time window between 2010 and 2020.

### Gradient forest modelling

We used the machine-learning gradient forest model to predict and rank the importance of these 1,088 weather metrics as predictors of CpG methylation levels, running 500 regression trees. The maximum number of splits was set to log_2_ (0.368 × number of predictor variables) / 2, as recommended by the package developer^79^. We considered weather metrics with scaled R^2^-weighted importance > 0.5 as having a significant effect, and these were used in subsequent analyses.

### Population genetic structure and the identification of weather-associated methylated CpG sites

The population genetic structure into which each individual was born was assessed to correct for latent factors in subsequent weather-association analyses. SNPs were called using BISCUIT 0.3.14^91^ and filtered using minor allele frequency for 0.1 and 20% missing data across samples using vcftools^92^. These SNPs were then pruned for linkage disequilibrium by purging sites with a pairwise *r*^2^ threshold of 0.1 with a window size of 50 Kb and a step size of 10 bp using PLINK v1.90. Eigenvectors and eigenvalues for principal component analysis (PCA) were also produced using PLINK v1.90. PCA was then used to evaluate methylation levels.

We used latent factor mixed modelling (LFMM) to test for any effect of the 33 significant weather metrics we detected on methylation levels. No prominent population genetic structure was detected, therefore *K* = 1 was used in LFMM. We ran LFMM with 3 repetitions with 1,000 maximum iterations and 500 burn-ins using the LEA 3.4.0 package. To control for bias and inflation in false positives, we first calculated the genomic inflation factor λ, defined as the observed median of *Z*-scores divided by the expected median of the chi-squared distribution for each environmental association, to calibrate *P-*values. We then corrected for multiple testing using the Benjamini and Hochberg method to obtain *Q*-values from adjusted *P*-values. Significant weather-associated CpG sites were determined as those with an absolute *Z*-score > 2 and a *q*-value < 0.05.

### Consequences of differences between adaptive and non-adaptive methylation on early-life weight

Using the 33 significant weather metrics as predictor variables, we again ran two gradient forest models, one using CpG sites not significantly associated with any weather metrics as response variables, and the other using only weather metrics associated with methylated (5mC) CpG sites as response variables. The PCAs of methylation turnover frequencies between these two models were compared using Procrustes rotation, where residuals implied the deviation of adaptive (i.e., weather-related) methylation from non-adaptive (i.e., weather-unrelated) methylation. We then tested the relationship between early-life weight at first capture and Procrustes residuals, as well as the relationship between cohort year and Procrustes residuals, using median-based linear model (MBLM).

### Epigenome-wide association analysis (EWAS) with early-life weight

The association between methylation levels and first recorded neonate weight was tested using latent factor mixed model (LFMM). The latent factor *K* was set at 1 and calibrations for genomic inflation were performed according to the same protocol used for LFMM analysis of weather-associated methylation. Ridge estimate was used to estimate the parameters of the LFMM model, based on minimising a regularised least-squares problem with an *L*_2_ penalty. Significant CpG sites were determined as those with a *Q*-value < 0.05.

### Promoter prediction and annotation of CpG sites

We defined promoter regions spanning 2,000 bp upstream to 200 bp downstream of a gene, for which CpG methylation could have a strong negative effect on gene expression, consistent with other animal studies^93–95^. Promoters were then annotated based on their proximity to the gene models in the mMelMel3.2 assembly. We used these gene models to search for homologous annotations in the genome of *Canis lupus familaris* (dog), canFam4 (GCF_011100685.1), which was the most closely related species in OrgDb database within the Bioconductor version 3.17 release.

Enrichment analysis was performed using clusterProfiler 4.0^96^ to detect over-representation of Kyoto Encyclopedia of Genes and Genomes (KEGG) pathways. Significant terms were determined as those with a *Q*-value < 0.05 after Benjamini-Hochberg correction.

## Results

### RRBS results and methylation calls

After trimming and filtering, we obtained an average of 4.34 M (± 1.96 M) paired-end clean reads for each sample. More than 90% of these reads had quality scores ≥ 30. These read data from Illumina sequencing for each sample were deposited in the NCBI Sequence Read Archive (SRR28700424–SRR28700518) under BioProject PRJNA1100394. After filtering, calling, and imputation for missing data, we obtained 37,672 autosomal CpG sites from these 95 badger blood samples.

### Gradient forest modelling for weather metric sliding windows

Among 1,088 weather metric sliding time windows, Pv_l04.l04 ranked top, followed by Tm_l03.l03, Tv_l06.l06, Pm_l04.l04, and Tv_l04.l04 (**Figure 2 and Supplementary Table 2**). While Pv was the most important metric for explaining the methylation pattern, the median frequency density of scaled R^2^-weighted importance was highest for Tm (∼0.7), followed by Tv, Pv, dTv, Pm, dPv, dPm, and dPv.

**Figure 2.**
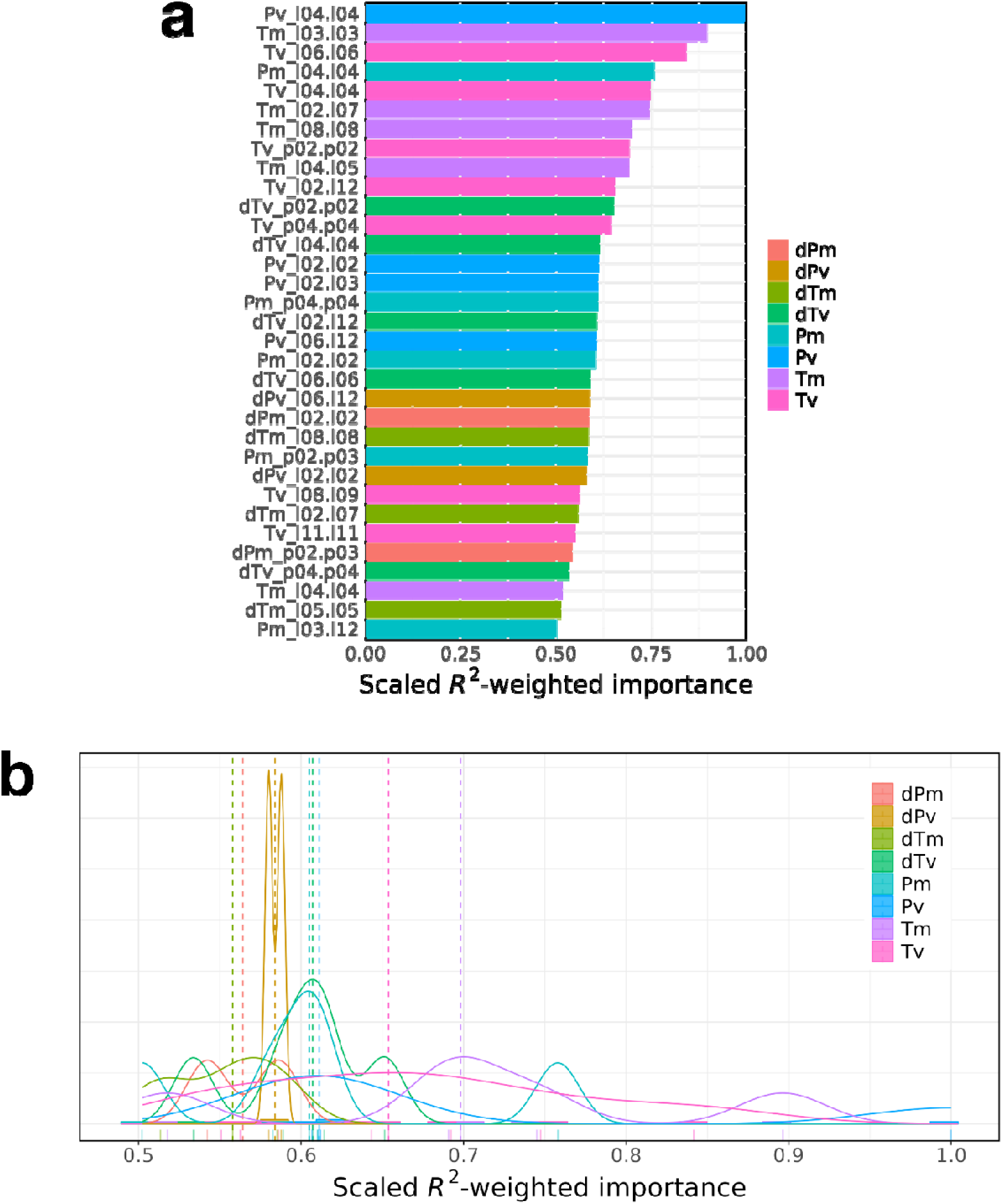
**(a)** Scaled *R*^2^-weighted importance of significant environment variables, only showing those > 0.5. **(b)** Frequency density plot of scaled R^2^-weighted importance of significant environment variable categories, only showing those > 0.5. Dotted lines show the median.

Each weather metric exhibited different time window phases of primary influence on methylation patterns (**Figure 1b**). For Pm (mean precipitation) and Pv (variation in precipitation), February and April of the previous year were most influential. For Tm (mean temperature), the greatest influence occurred between March and May of the birth year, as well as during February and July in the previous year. For Tv (variation in temperature), effects were greatest in April and June of the birth year, with some effect attributable to cohort (i.e., year of birth). We found no significant importance hotspots for dPm (deviation of mean precipitation from 10-year average), dPv (deviation of variation in precipitation from 10-year average), dTm (deviation of mean temperature from 10-year average), and dTv (deviation of variation in temperature from 10-year average), there were no significant importance hotspots. These generally also had a lower level of importance than Pm, Pv, Tm, and Tv.

Most of these important time windows occurred along the diagonal depicted in the heatmap (**Figure 1b**). This implies that variations in weather metrics over short time-windows were more important in explaining methylation pattern than longer-term variations.

### Weather effect-associated CpG sites

Principal component analysis revealed no prominent genetic structure among cohorts (**Figure 3**). LFMM analysis detected 4,641 CpG sites (12.32%) significantly associated with at least one significant weather metric (|Z| > 2 and *Q* < 0.05), after correcting for genomic inflation and multiple testing (**Supplementary Table 3 and 4**).

**Figure 3.**
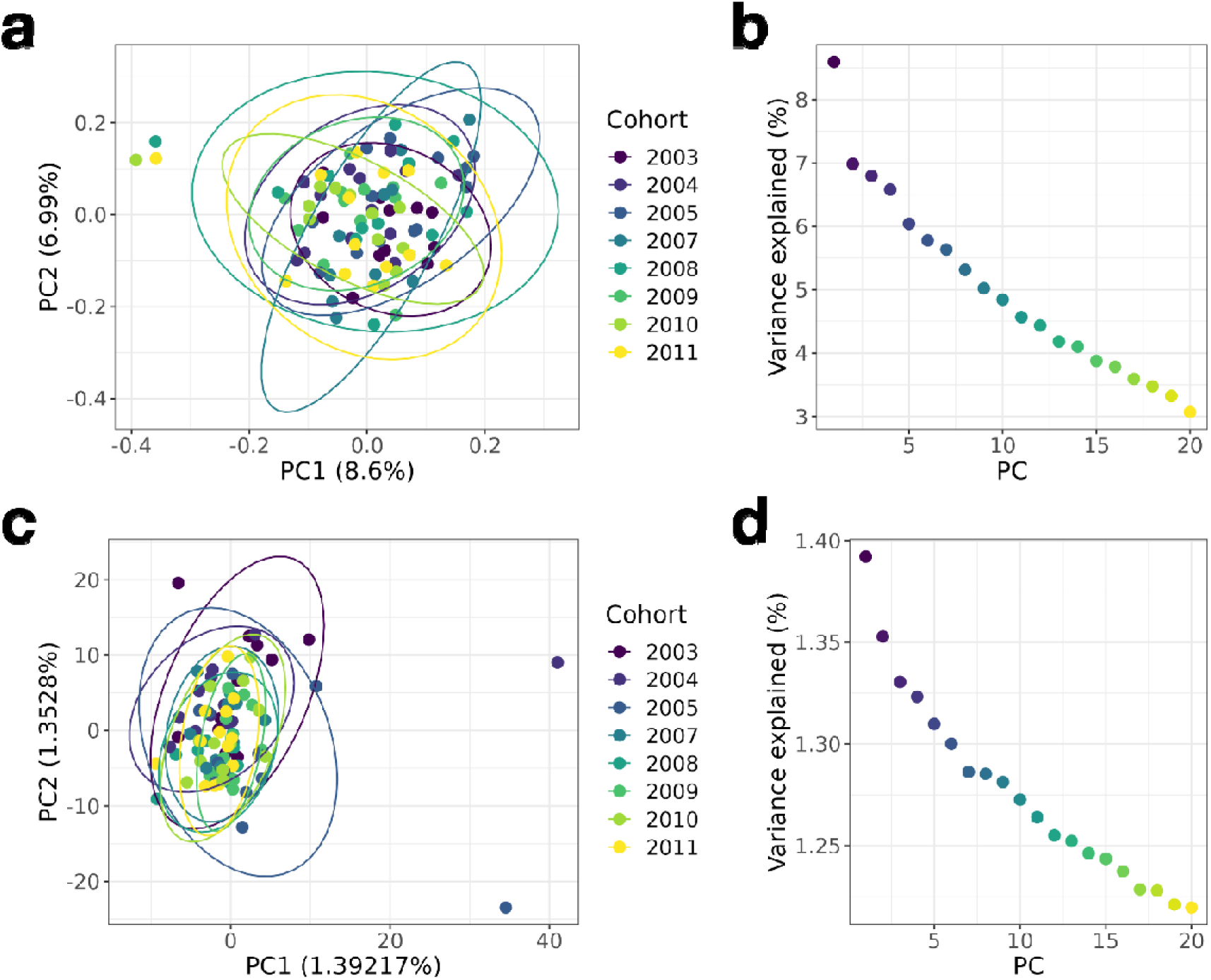
**(a)** Genetic structure of cohorts visualised from the first two principal axes of the principal component analysis (PCA), with **(b)** the variance explained by 20 principal axes.

Only four of these significant weather-associated CpG sites were situated in gene bodies, while none occurred in promoter regions (**Supplementary Table 5**). One CpG site, chr10_224521, was in the *GAS2L1* gene (growth arrest specific 2 like 1; ENSSEMG00010017771) and significantly associated with dTv_p02.p02. Two CpG sites, chr10_249503 and chr10_249521, were in the *CACNA1B* gene (calcium voltage-gated channel subunit alpha 1B; ENSSEMG00010017084) and significantly associated with Tm_l02.l07, Tm_l04.l05, dTm_l02.l07, dTv_l04.l04, and dTv_p02.p02. One CpG site, chr3_2174752, was significantly associated with Tv_l08.l09 and dTm_l02.l07, but the function of its corresponding gene is unknown.

Another 14 sites were located in the 47S rDNA region (5’ETS–18S–ITS1–5.8S– ITS2–28S–3’ETS) on chromosome 16 (**Figure 4a and b; Supplementary Table 5**). Many of these 47S rDNA CpG sites were concurrently associated with more than one weather metric. In particular, five CpG sites (i.e. chr16_61045, chr16_61073, chr16_61083, chr16_61089, chr16_61123), which were all situated in the 18S intragenic region, showed significant association between their methylation level and dPv_06.12 (deviation of variation in precipitation between June and December to 10-year average) (**Figure 4c**).

**Figure 4.**
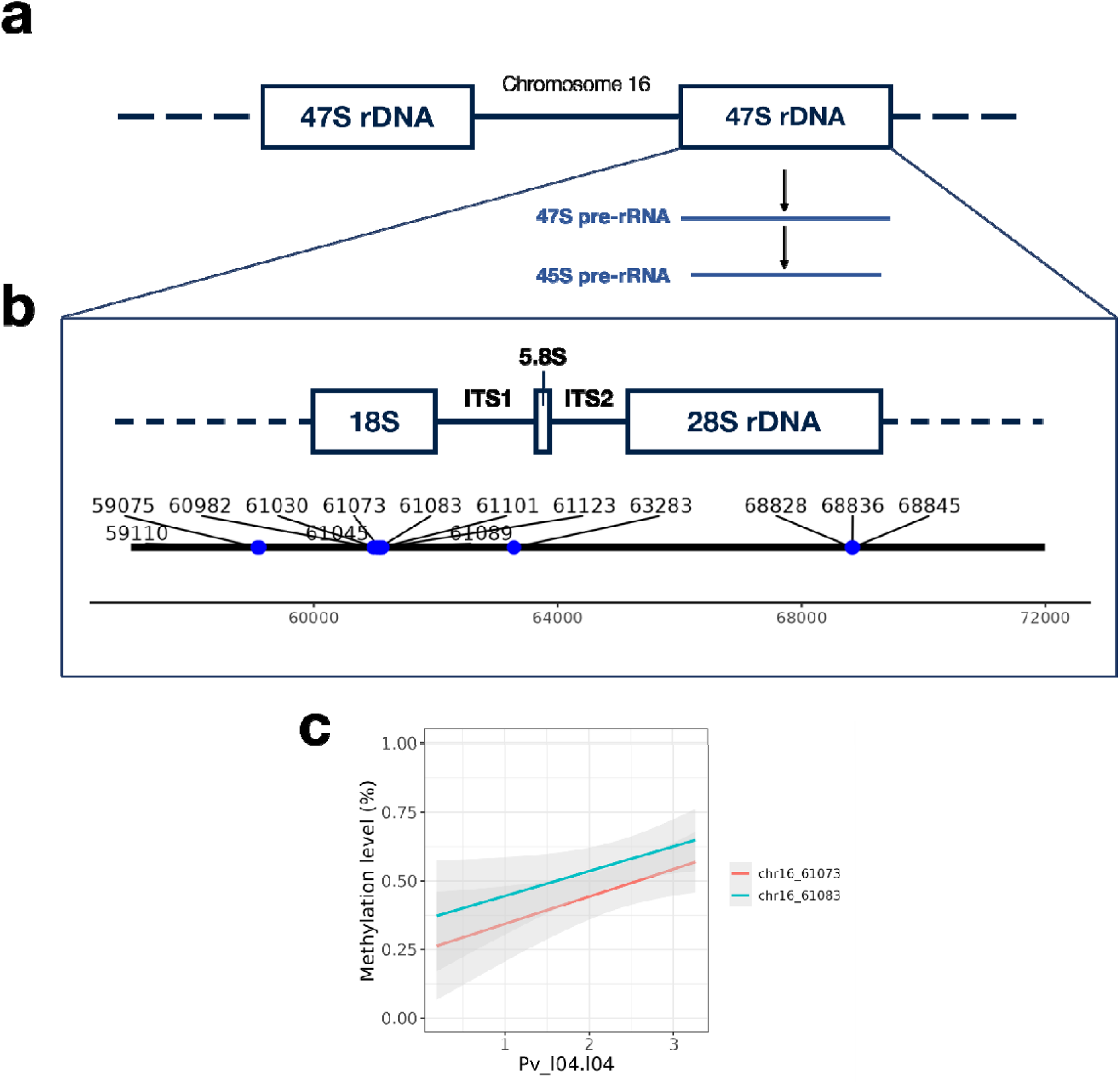
**(a)** Tandem-repeated 47S rDNA cluster on chromosome 16 of the *M. meles* genome. **(b)** Structure of a 47S rDNA cluster. Blue dots along the number line denote significant environment-associated methylated CpG site. **(c)** Association between methylation level and the environmental variable Pv_l04.04 for two CpG sites within the 47S rDNA cluster. **(d)** Association between methylation level and the environmental variable dPv_l06.l12 for five CpG sites with the 47S rDNA cluster.

### Procrustes residuals

Linear regression between early-life weight and cohort size revealed no significant trend (*r* = 0.15, *P* = 0.1418). The coefficient of effect of cohort size was only 0.0039 (**Supplementary** Figure 1). This ruled out the potential effect of negative density-dependence, which might confound the subsequent analyses with early-life weight.

Early-life weight increased significantly with Procrustes residuals (*P* = 5.07e–07) by a coefficient estimate of 0.73 (**Figure 5a**). In contrast, we did not detect any significant association between Procrustes residuals and time (*P* = 0.266) and the coefficient estimate of the effect of time was very small, only 0.010 (**Figure 5b**).

**Figure 5.**
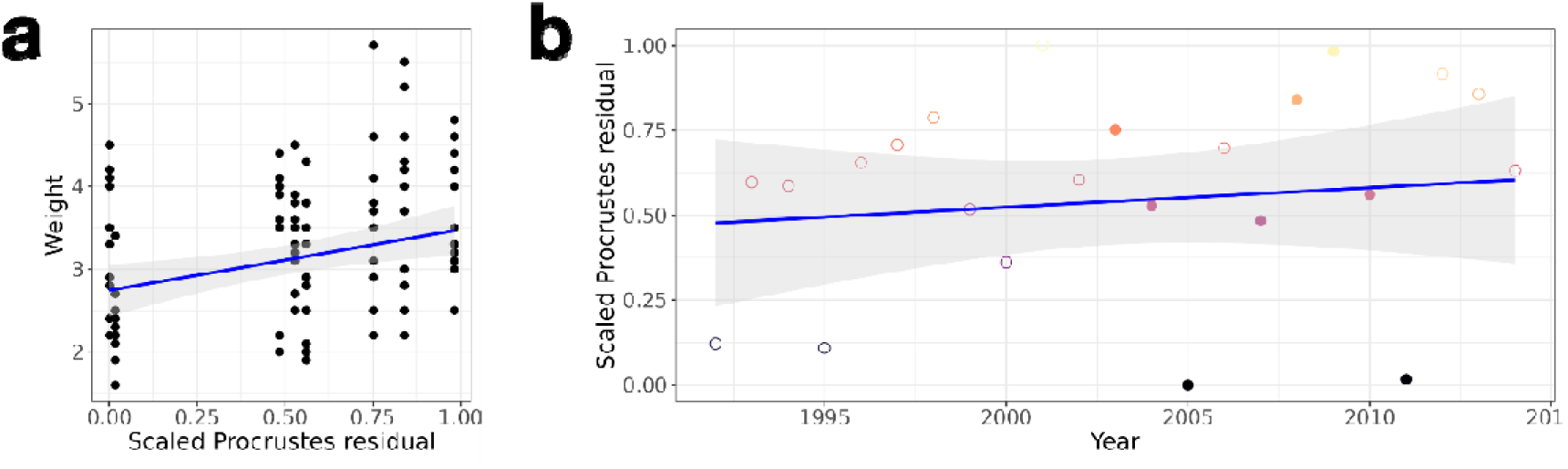
**(a)** Association between cub weight at three to four months old and min-max scaled Procrustes residual. **(b)** Scaled Procrustes residual of the cohorts across time. The solid dots represent those included in this study. The hollow dots represent those not included in this study but predicted based on long-term weather data.

### Early-life weight-associated CpG sites

LFMM analysis detected 1,641 CpG sites (4.36%) that were significantly associated with early-life weight (*Q* < 0.05) (**Figure 6a**). There were 32 CpG sites in potential promoter regions and 600 CpG sites in gene bodies of the 525 gene models for badgers (**Supplementary Tables 6 and 7**). Three KEGG pathways were over-represented in these promoter or gene body CpG sites (*Q* < 0.05; **Supplementary Table 8**), namely cfa04072 (phospholipase D signalling pathway; **Supplementary** Figure 2), cfa04727 (GABAergic synapse), and cfa05032 (morphine addiction).

**Figure 6.**
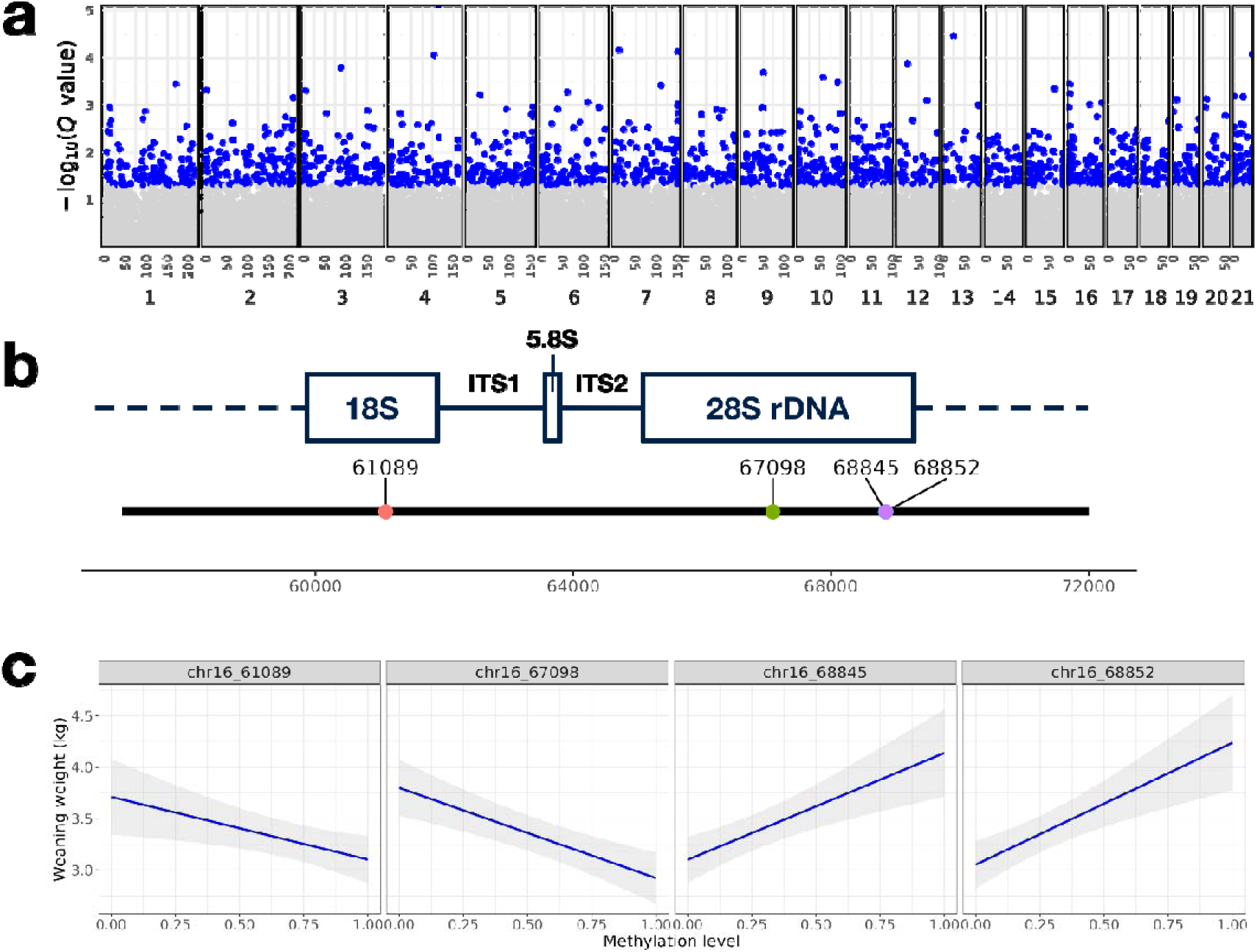
**(a)** Manhattan plot of epigenome-wide association with badger early-life weight. Blue dots show significant CpG sites (*Q* < 0.05 and grey dots show insignificant sites (*Q* > 0.05). **(b)** The locality of the four significant CpG sites within the 47S rDNA cluster. **(c)** Association between methylation level and weaning weight for the four significant CpG sites.

Within the 47S rDNA, we detected 4 significant CpG sites (i.e., chr16_61089 within the 18S portion, and chr16_67098, chr16_68845, and chr16_68852 within the 28S portion) (**Figure 6b**). The methylation levels of chr16_61089 and chr16_67098 were negatively correlated with early-life weight, while that of chr16_68845 and chr16_68852 were positively correlated with weight (**Figure 6c**).

## Discussion

### Methylation-weather association reveals type and timing of selection pressures

Variability in precipitation (Pv) in the early spring during pre-implantation was the top determinant of methylation patterns in neonatal badgers, followed by mean temperature (Tm) and temperature variability (Tv), also in spring months. This is congruent with broader understanding of seasonal weather effects on badgers, where spring weather conditions that are either too warm or cold, or too wet or dry, as well as substantial variation from mean precipitation during early summer^50,52,86^, are associated with lower badger survival rate, impaired cub growth rates^97^, and lower reproductive success^47,49,53,54,98^. Furthermore, these effects are exacerbated by higher coccidiosis-related mortality rates among badger cubs in poor weather years^54^

Although lower autumn rainfall also presents a risk factor associated with poorer badger body condition in winter^50,52^, autumnal windows did not affect methylation patterns in this study population. This implies that *in utero* epigenetic programming may be affected by environmental stimuli more intensively in the earlier (spring) stages of delayed implantation.

Long-duration delayed implantation occurs in >100 species of mammals^61,99,100^, with blastocysts undergoing dynamic metabolic changes during diapause^61,67–69^. In mice, previous work has found a slight but statistically significant increase in DNA methylation level at one locus in diapause blastocysts via experimentally delayed implantation compared to 4.5-embryonic-day blastocysts without manipulation^101^. Similarly, maternal hormonal changes result in differential methylation patterns between dormant and implanted (E2-activated) mice blastocysts^75^. Therefore, *de novo* methylation during delayed implantation can play a significant role in adaptative developmental programming, as implied by the effects we found in 4 CpG sites in genic regions and in 14 sites located in 47S rDNA.

### 47S rDNA can translate environmental stress into phenotypic stress

In eukaryotes, the ribosome is responsible for cellular protein synthesis, with translation encoded by ribosomal RNA (rRNA) organised on chromosomes in tandem-arrayed clusters^102^. rDNA, transcription units are separated by large, intergenic non-transcribed spacers (IGS) within these clusters^103^. In mammals, these rDNA clusters are 47S and 5S. In badgers, 5S clusters are distributed across all chromosomes, however 47S clusters are only found on chromosomes 16 and Y. During eukaryotic ribosome biogenesis, part of the 5’ETS and all of the 3’ETS of 47S rRNA is initially cleaved to form 45S rRNA, and subsequently to yield 18S, 5.8S, and 28S rRNA^104^. All mature rRNAs are then assembled into ribosomal subunits.

rDNA can be activated or inhibited by epigenetic mechanisms^105^. In badgers, most of the significant weather-associated CpG sites occurred in the 18S and ITS1 regions, within chr16, and thus without male bias relating to chrY. The methylation rate of chr16_61073 and chr16_61083, located in the 18S gene body, increased with variation in precipitation.

Hypermethylation was associated with heavier weaning weight in badgers. Poor maternal diet during the preimplantation period in lab mice alters rDNA methylation and downregulates rDNA expression, reducing cellular RNA content in the foetal somatic tissues, which only recover after diet restriction is remediated^106^.

Notably, out of thousands of CpG sites we discovered, most that had a significantly weather-association were located on rDNA. This further supports that rDNA appears to be a critical but rare methylomic target during prenatal development ^107^. Given that neonatal badger cubs suffer impaired nutrition and growth due to coccidia infection^55^, leading to > 60% mortality in cohorts during dry years^56^, future research should investigate whether maternal malnutrition triggers rDNA methylation promoting genes linked to greater coccidiosis resistance or lower coccidiosis morbidity^108,109^.

### Prenatal methylation change alters postnatal fitness

The positive association between scaled Procrustes residuals and early-life weight implies that environmental conditions affecting prenatal development *in utero* caused adaptive postnatal phenotypic changes. The larger the Procrustes residuals, the stronger the deviation between adaptive-environment association and neutral-environment association, suggesting a pattern of local adaptation^110,111^. This supports that epigenetic changes in badgers increase the likelihood that the individual’s phenotype will be better matched to environmental conditions (i.e., a predictive adaptive response, PAR^112^), enhancing its probability of survival and thus ultimately its fitness, irrespective of a constant, fixed genotype^113,114^. Of note, we found no significant association between scaled Procrustes residuals and time; that is, methylation change in badgers did not appear to follow any temporal trend.

Of the 1,641 CpG sites for which differential methylation affected early-life weight, only chr16_61089 and chr16_68845 overlapped with those associated with weather stress. These two rDNA loci thus play a pivotal role in translating environmental stress into phenotypic response via epigenetic status, although they function antagonistically, such that the methylation level of chr16_61089 was negatively associated with early-life weight, while that of chr16_68845 was positively associated. This highlights the heterogeneous effects of gene body methylation under various conditions and warrants further mechanistic investigation^115,116^.

### Phospholipase D signalling and GABAeric synapse

Three KEGG pathways were over-represented in genes for which differential methylation was associated with early-life weight, namely cfa04072 (phospholipase D signalling pathway; **Supplementary** Figure 2), cfa04727 (GABAergic synapse), and cfa05032 (morphine addiction).

In mammals, phospholipase D (PLD) is central to various cellular signalling components^117^. The phospholipase D signalling pathway also involves the hydrolysis of phosphatidylcholine by PLD to produce phosphatidic acid (PA), which acts as a secondary messenger in a series of signalling cascades^118^. We identified that many key genes in this pathway were differentially methylated and associated with variation in early-life weight: Phosphoinositide 3-kinases (PI3K) regulate cell growth, motility, survival, and metabolism^119^; GPCR regulates PLD activity^120^; Protein kinase C (PKC) phosphorylates and stimulates PLD^121^. Most importantly, the crosslinking of the high-affinity IgE receptor FcεRI via IgE-antigen complex far upstream of this pathway has implications in inflammatory responses ^122^. This suggests a mechanism for early-life epigenetic responses to regulate immunity and pathogen defence, potentially helping badger cubs survive severe juvenile coccidiosis^55^. However, the development and maintenance of an effective, pre-adaptive immune system could be costly and has been seen to result in a trade-off between growth and immune function in animals^123,124^, as well as risking later life autoimmune diseases or pleiotropic carcinomas.

Regarding cfa04727 (GABAergic synapse), we identified that the central GABA-A and GABA-B receptors, which mediate inhibitory neurotransmission^125^, and GABA transaminase (GABA-T), which converts GABA to succinic semialdehyde^126^, were methylation targets. In relation to early life-history stages, GABA synapses regulate the maturation and activity of neural networks during brain development, controlling neuronal migration, synapse formation, and synchronisation of neural network activity^127^. Experience-dependent processes and activity-driven mechanisms are essential for the development of GABAergic circuits. Therefore, methylation of genes in this pathway could potentially underpin synaptic plasticity, which regulates network activity in the brain in response to varying environmental constraints^128^. Particularly, GABA interacts with dopamine and serotonin to modulate aggressive behaviour^129^, and the GABAergic neuronal system mediates the gut-vagus-brain pathway^130^. Therefore, weather-related metabolic stress and/or a scarcity of food may promote *in utero* or post-natal methylation that could influence how aggressive and competitive badgers are during times of weather-driven food constraint, where badgers can be extremely fierce^131^.

### From lab to field: environmental epigenetics and climate change adaptation

Our study presents the first evidence for prenatal epigenetic change in response to environmental conditions in a wild animal. This represents an important milestone moving from laboratory studies on epigenetic effects to examine their broader ecological and evolutionary consequences in nature. Importantly, we established that badgers utilise methylation as a means to adapt developmental programming during the preimplantation blastocyst stage, with broader implications for other organisms that have evolved delayed implantation, as well as for embryonic development in general. Ribosomal DNA provides a major mechanism for relaying environmental stress into phenotypic effects during early-life development, with implications for subsequent fitness. Genotypic adaptation, via Darwinian selection, takes generations to occur^132^, which is particularly challenging for species with longer inter-generational times^133^. While badgers exhibit complex mate choice behaviours, selecting for inbreeding avoidance^63^, heterozygosity^134^, and immunogenetic fitness^135^, the instantaneous effects of epigenetic methylation on the expression of key genes may enable immediate adaptation to weather and food supply conditions anticipated in the birth year, potentially operating through enhancing an individual’s resistance to endemic juvenile disease^54^.

Schloss et al.^136^ predicted that, for the western hemisphere, an average of 9.2% of mammal species at any given location will be unable to respond to climate change. This was, however, based on intergenerational rates of selective adaptation, where our work suggests that it may not be appropriate to assume changes from current normative climatic conditions will so readily exceed the epigenetic adaptability of species.

## Author contributions

T.H.H.: designed the study, conducted the experiment and bioinformatic analyses, and drafted the manuscript;

M.-S.T.: designed the study, collected and processed the samples, and drafted the manuscript;

C.N.: collected the samples and revised the manuscript;

D.W.M.: collected the samples and revised the manuscript;

C.D.B.: collected the samples and revised the manuscript.

