## Supplementary Information for "Pre-implantation genome-wide methylation enables environmental adaptation in a social meso-carnivore"

Tin Hang Hung^1,*,^^
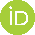
<https://orcid.org/0000-0001-9853-2053>, Ming-shan Tsai^2,^^
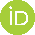
<https://orcid.org/0000-0001-6804-1317>, Chris Newman^2^
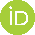
<https://orcid.org/0000-0002-9284-6526>, David W. Macdonald^2^
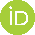
<https://orcid.org/0000-0003-0607-9373>, Christina D. Buesching^3^
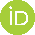
^<https://orcid.org/0000-0002-4207-5196>^

1. Department of Biology, University of Oxford, OX1 3RB, United Kingdom
2. Wildlife Conservation Research Unit, The Recanati-Kaplan Centre, Department of Biology, University of Oxford, OX13 5QL, United Kingdom
3. Department of Biology, Irving K. Barber Faculty of Sciences, The University of British Columbia, Okanagan, Kelowna, British Columbia, Canada

^T.H.H. and M.-S.T. contributed equally as co-first authors.

Corresponding author:

*T.H.H.:

**Supplemental Table 1.** Count of samples selected from each cohort from 2003 to 2011 (the samples of 2006 were not collected) for this study. All protocols and procedures were approved by the Animal Welfare and Ethical Review Board, Department of Zoology, University of Oxford and were conducted under the Animals (Scientific Procedures) Act 1986 (PPL: 30/3379)

| **Cohort** | **Male** | **Female** | **Total** |
| --- | --- | --- | --- |
| 2003 | 4 | 7 | 11 |
| 2004 | 6 | 7 | 13 |
| 2005 | 6 | 7 | 13 |
| 2007 | 6 | 3 | 8 |
| 2008 | 5 | 7 | 12 |
| 2009 | 6 | 6 | 12 |
| 2010 | 5 | 7 | 12 |
| 2011 | 6 | 7 | 12 |
| Total | 43 | 50 | 95 |

**Supplementary Table 2.** Scaled *R*^2^-weighted importance of all 1,088 environmental variables, arranged in descending order of importance

*See separate spreadsheet*

**Supplementary Table 3.** *Z*-scores of association between the methylation level of CpG sites and 33 significant environmental variables

*See separate spreadsheet*

**Supplementary Table 4.** *Q*-scores of association between the methylation level of CpG sites and 33 significant environmental variables

*See separate spreadsheet*

**Supplementary Table 5.** Annotation of significant methylation-weather-associated CpG sites

*See separate spreadsheet*

**Supplementary Table 6.** Annotation of significant methylation-weight associated CpG sites in potential promoter regions

*See separate spreadsheet*

**Supplementary Table 7.** Annotation of significant methylation-weight associated CpG sites in gene bodies

*See separate spreadsheet*

**Supplementary Table 8.** Results of Kyoto Encyclopedia of Genes and Genomes enrichment analysis on the methylation-weight associated CpG sites in promoters or gene bodies

*See separate spreadsheet*

**Supplementary Figure 1.** Enriched KEGG pathways over-represented in genes that contain significant weight-associated CpG sites, including **(a)** the phospholipase D signaling pathway (cfa04072), **(b)** the GABAergic synapse (cfa04727), **(c)** and the morphine addiction (cfa05032). Data are from KEGG graph and rendered by Pathview.

**a**

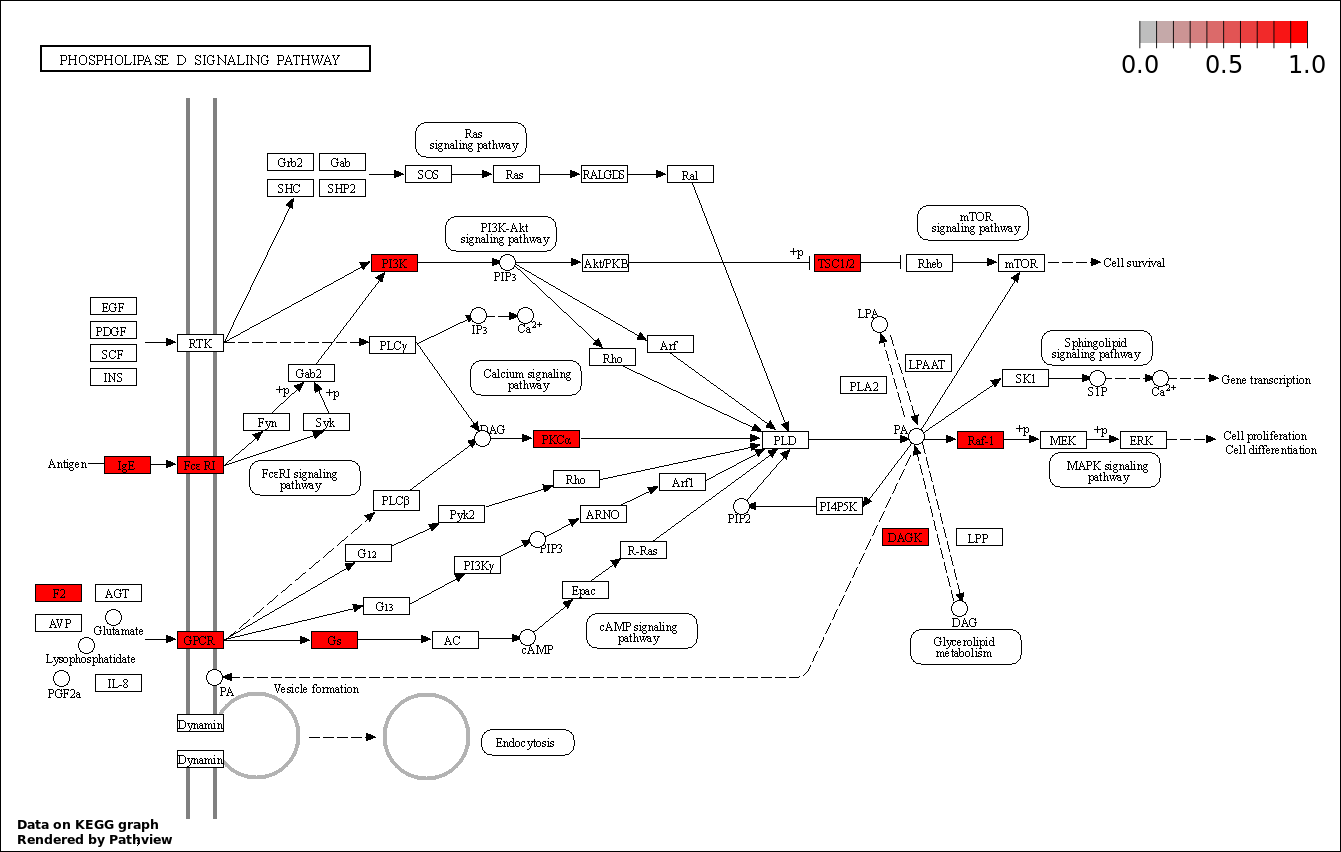

**b**

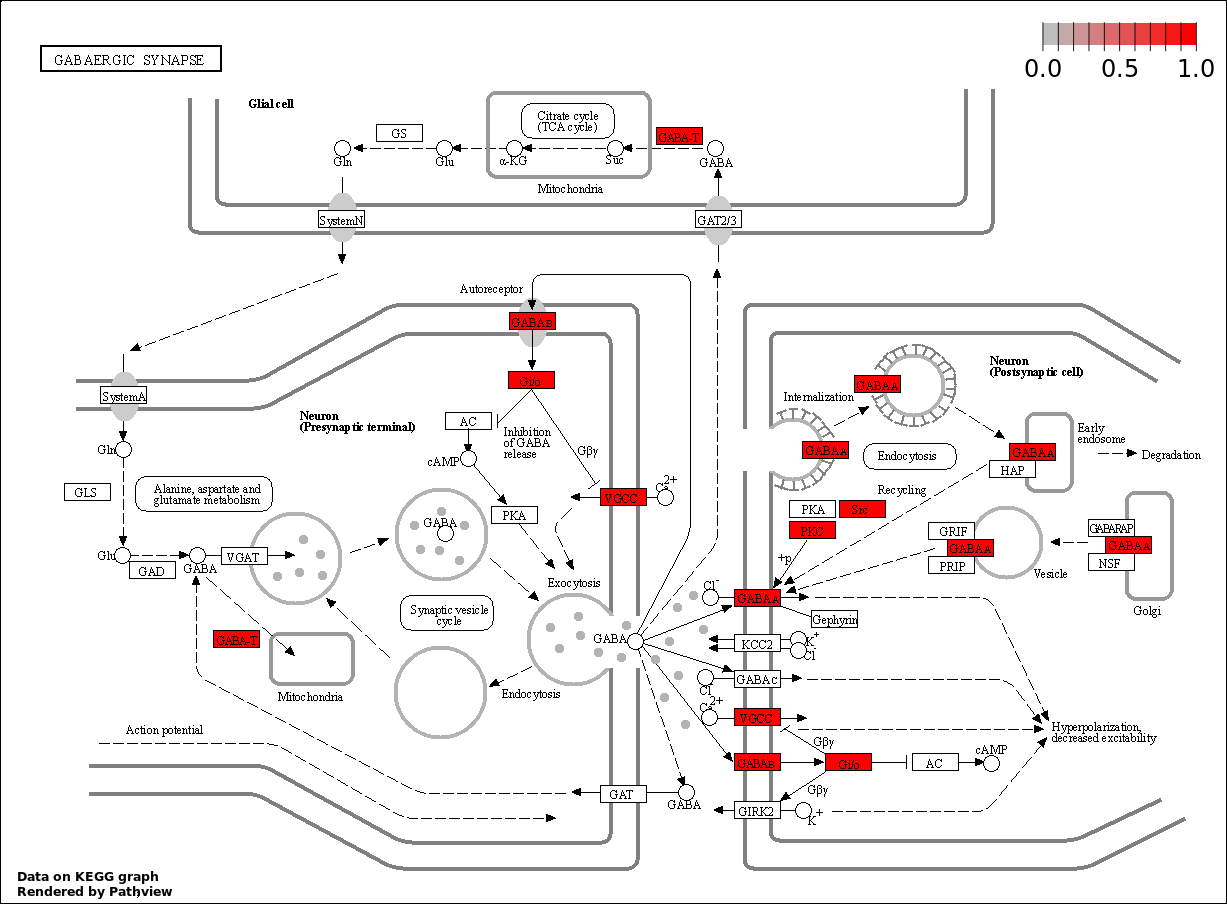

**c**

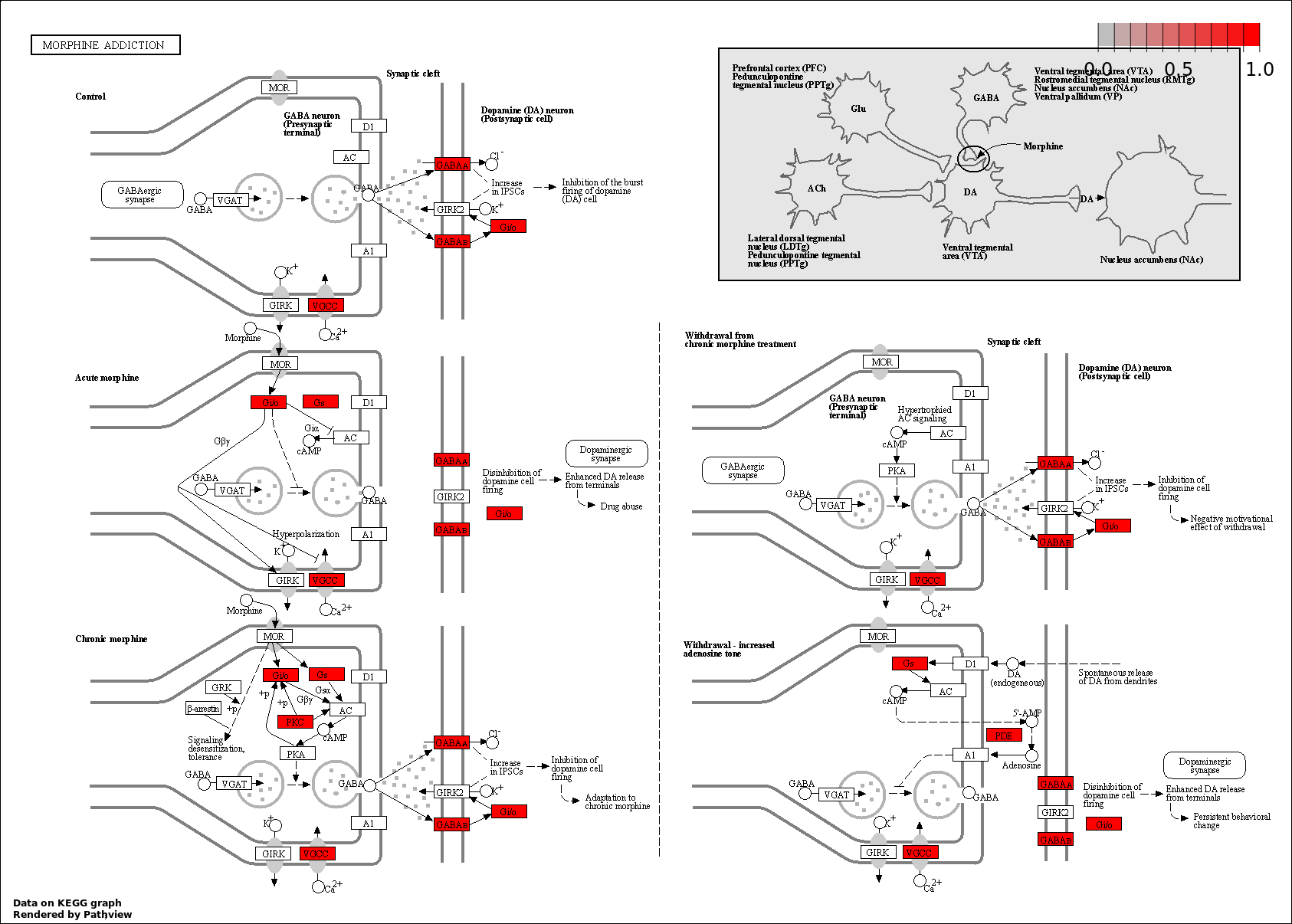
